# Hypoxia can set the elevational limits of dragonflies

**DOI:** 10.64898/2026.10.13.758859

**Authors:** Michael P. Moore, Anna Brasket, Megan Mokriski, Karla Alvarez, Tania Steinmeier

## Abstract

Oxygen declines by ∼10% for each 1000m gained above sea level. Although the impact of low oxygen on endotherms has received considerable attention, we know far less about whether or not hypoxia can set the elevational limits of terrestrial ectotherms. Here, we addressed this question using Nearctic dragonflies. First, we experimentally tested if dragonflies are vulnerable to the low oxygen found at high elevations. Our results show that even resting adults struggle with the P_O2_ that occurs at the highest elevations where dragonflies are found in Colorado, USA. Second, we used a comparative approach to test if the dragonfly species that can inhabit higher elevations are also more tolerant of low oxygen. Our analysis shows that a species’ upper elevational limit is precisely predicted by its vulnerability to low oxygen. Collectively, these results provide some of the first evidence that low oxygen can be an important constraint on the elevational maxima of terrestrial ectotherms.

## INTRODUCTION

Animals encounter many harsh conditions at high elevations [1,2], but one of the most serious is low oxygen [2–4]. Insufficient oxygen relative to physiological demands (“hypoxia”) can inhibit essential activities, like foraging and mating, or even disrupt the basic requirements of life, like digestion and brain function [3,5,6]. Thus, to live high in the mountains, species must be able to tolerate the low oxygen [3]. Nevertheless, the impact of hypoxia on species’ upper limits is only known for a handful of endotherms (e.g. [5,7,8]). By contrast, despite representing the vast majority of the world’s animal diversity, very little is known about whether or not hypoxia shapes the upper limits of terrestrial ectotherms [3].

To date, the risk of hypoxia for high-elevation ectotherms has been somewhat contentious. Some have argued that hypoxia is unlikely to be a major issue for terrestrial ectotherms in the mountains (e.g. [9]). Since ectotherms do not metabolically maintain their own internal temperatures, their relatively low resting demands for oxygen might never exceed the supply of oxygen at high elevations [6,10,11]. Similarly, low oxygen may only be an issue for ectotherms when they are hot and metabolic demands are very high [12–14]—a problem that might never occur in the cooler montane climates. Despite these arguments and others, recent empirical studies have also implied that low oxygen could be a more significant problem for high-elevation ectotherms than is presently understood [3,15,16]. For instance, insect species with the highest demands for oxygen are dispersing to higher elevations more slowly than those with lower demands [17]. Seemingly in response to the low-oxygen conditions, some high-elevation ectotherms even have adaptations that should improve the delivery and use of oxygen [18]. Given this conflict between the physiological calculations and the indirect empirical evidence, we need studies that directly test if low oxygen could set species’ elevational limits.

Here, we used dragonflies to address two key questions about hypoxia’s impact on the elevational limits of ectotherms. First, is the low oxygen found at high elevations a general problem or only a problem at high temperatures? To address this question, we manipulated both oxygen and temperature and then tested if low oxygen kills adult dragonflies faster or only faster at high temperatures. Second, are those species that inhabit high elevations more tolerant of low oxygen? For this question, we estimated mortality rates under low oxygen for 9 different species and tested if their tolerances predicted their upper elevational limits in the Rocky Mountains.

## METHODS

### Study System and Collections

Dragonflies are very well-suited to studying the challenges posed by low oxygen for several reasons. First, dragonflies have extraordinary metabolic demands for oxygen [19,20]. Second, dragonflies have established in most biomes across the planet, yet very few species live >3000m asl, and no species have been recorded consistently living >4000m asl [21]. Third, recent comparative analyses indicate that the species that do live at high elevations have biomechanical advantages that reduce the aerobic costs of flight [22]. Thus, if any insect is likely to be sensitive to the low-oxygen conditions at high elevations, dragonflies are a good candidate.

We field collected 164 adult males from one of 3 ponds all in Denver, USA below 1680m asl. Upon capture, we brought adults to our lab at the University of Colorado Denver and held them overnight in loosely covered plastic petri dishes at 4ºC. The next day, we moved the dished dragonflies to an incubator set to 18ºC and warmed them up prior to the experimental trials.

### Is low oxygen a general problem for dragonflies or only a problem at high temperatures?

We began by conducting a 2x2 factorial experiment, manipulating temperature and oxygen, and measuring how long each adult survived. In this experiment, we placed dragonflies in their dishes into a DigiTherm^®^ Incubator set to one of two temperatures and one of two oxygen levels. For our temperature treatments, we used either a near-optimal temperature for performance (39ºC) or a hot but sub-lethal temperature that dragonflies reach in the wild when defending territories (47ºC). These conditions were based on prior estimates of the thermal performance curves of dragonflies [23,24]. For our oxygen treatments, we used either the ambient oxygen conditions for our study location (P_O2_=130.7mm Hg, equivalent to ∼1600m asl) or the low-oxygen conditions that occur ∼150m above the highest record of dragonflies in Colorado, USA (P_O2_=95.7mm Hg, equivalent to ∼4,100m asl). To manipulate the oxygen levels inside the incubator, we used a Hypoxico^®^ Sierra 100 altitude generator, which filters O_2_ out of the ambient air and flows a specified gas mixture into a space without altering overall air density. We flowed air into the incubator at 100mL/min by inserting the altitude generator’s outflow hose into a hole that was drilled through the sidewall of the incubator [11]. All manipulations were performed in the same incubator on different days. We performed one experimental treatment per day, and so all dragonflies that were captured on the same day were exposed to the same combination conditions (e.g. high temperature/low oxygen or low temperature/high oxygen; see also [11] for validation of these methods). We performed this experiment on 124 individuals from 3 species that are common to low-elevation ponds in Colorado but have differing upper elevational limits: *Pachydiplax longipennis* (∼1800m asl), *Erythemis simplicicollis* (∼2500m asl), and *Libellula pulchella* (∼3300m asl).

After placing dragonflies in the incubator, we checked if each adult was alive every 20 minutes for 2 hours. To check the status of each dragonfly, we first examined its dish for any movement. If no movement was observed, we gently shook the dish to check if the dragonfly showed any sign of life. If the dragonfly still did not move, it was considered dead and moved to an environmental chamber set to 18ºC. No dragonflies ever woke up after being scored as dead. We attempted to film these trials using a GoPro^®^ Hero 11, but the camera consistently overheated in the incubator and shut itself off. However, an earlier study demonstrated that opening the door for a short period of time does not affect the oxygen levels within the dishes (see [11]). Each individual’s time of death was considered the checkpoint of the trial when it was observed dead (i.e. the end of the 20-minute interval). Dragonflies that lasted the full 2 hours were scored as surviving.

To compare mortality rates among the four treatments, we fit mixed-effects Cox Proportional Hazards Models using the R package ‘coxme’ [25]. Adults that died during the treatment were scored as undergoing the “event” (event=1), with their death time as the time of their last known fate. Adults that did not die during their trial were scored as not having an event (event=0), with 120 minutes as the time of their final known fate. We fit a full model that included fixed effects for oxygen treatment, temperature treatment, and their interactions. We also fit separate models for each simpler iteration of fixed effects (Table S1). To account for any non-independence of adults that were from different species, or that were tested in the same treatment but on different days, we included random intercepts for species and date. We competed models against each other using an AIC_C_ model comparison approach, and we report the fold-change in mortality rate induced by the treatments±SE and 95% CIs. If low oxygen is a consistent problem for dragonflies, then the best-supported models should include a main effect of oxygen, where mortality rates are higher in the low-oxygen treatments. By contrast, if low oxygen is only a problem at high temperatures, then the best-supported model should include the interaction between the oxygen and temperature treatments, and low oxygen should have a greater impact at high temperatures than at normal temperatures.

### Does a species’ sensitivity to low oxygen conditions shape its upper elevational limit?

We next tested if the species that have higher elevational limits in Colorado are less vulnerable to low oxygen. For this test, we used 109 adults from 9 species and measured their mortality rates at 47ºC under either ambient vs low oxygen (see Supporting Information). Specifically, we targeted species from a single family that are common at low elevations but have dramatically different upper elevational limits: *Pachydiplax longipennis* (∼1800m asl), *Perithemis tenera* (∼1700m asl), *Libellula pulchella* (∼3300m asl), *Libellula luctuosa* (∼2100m asl), *Libellula forensis* (∼2600m asl), *Erythemis simplicicollis* (∼2500m asl), *Sympetrum pallipes* (∼3000m asl), *Sympetrum obtrusum* (∼2900m asl), and *Sympetrum corruptum* (∼3900m asl). To address our specific question, we focused on the mortality rates of 57 adults that were tested under low oxygen, and we report the full experimental results across both treatments in the Supporting Information.

We then generated estimates for species’ vulnerability to low oxygen and their current upper elevational limit in Colorado. To estimate each species’ vulnerability to low oxygen, we fit a Cox proportional hazards model to all individuals in the low-oxygen/high temperature treatment combination. This model included species as the only predictor, which allowed us to estimate the hazard for each species at each 20-minute checkpoint. Species had a median±SD of 5±5 (range:1-19) replicates in this treatment. As our estimate for each species’ vulnerability to low oxygen, we used the model-estimated cumulative hazard at the final check point. Next, to estimate each species’ upper elevational limit in the Rocky Mountains, we used the report of a survey of the state of Colorado, USA by Prather & Prather [26] and the maximum GBIF record for each species from the state [27]. The estimates from these two sets of records were typically similar, and so we used the higher maximum between the two.

To test if dragonfly species with higher elevational limits are less sensitive to low oxygen, we used phylogenetic generalized least squares [28]. We included a species’ model-estimated cumulative hazard rate as the predictor variable and its upper elevational limit as the response variable. To improve the normality of residuals, we ln-transformed both variables. To account for the shared evolutionary history among species, we included the phylogeny produced by Rocha-Ortega and colleagues [29]. We used AIC_C_ to compare models that assumed a Brownian Motion model of evolution, a Pagel’s λ branch-length transformation, and a star phylogeny. This approach indicated that a star phylogeny was the best fit (Table S2), and we therefore report results from this model. In this approach, if a species’ vulnerability to low oxygen is linked to its upper elevational limit, then we should find a negative relationship between a species’ cumulative hazard rate and its observed elevational maxima.

## RESULTS

### Is low oxygen a general problem for dragonflies or only a problem at high temperatures?

The best-supported model of mortality included only the main effects for temperature treatment and oxygen treatment (w=0.615). In this model, higher temperatures increased mortality rates by 10.7x±0.4 (95%CIs: 4.9 to 23.6), and low oxygen increased mortality rates by 3.4.x±0.3 (95%CIs: 1.8 to 6.4). This top model without the interaction was 1.7 times more likely than the model with the interaction (w=0.357), and it was >27 times better than every other model (Table S1). Even in the model with the interaction, however, the effect of low oxygen was not notably greater at high temperatures (95%CIs of estimate: 1.8x to 7.7x; Fig. 1A) than at normal temperatures (95%CIs of estimate: 95%CIs: 0.7x to 9.6x; Fig 1B).

**Fig 1.**
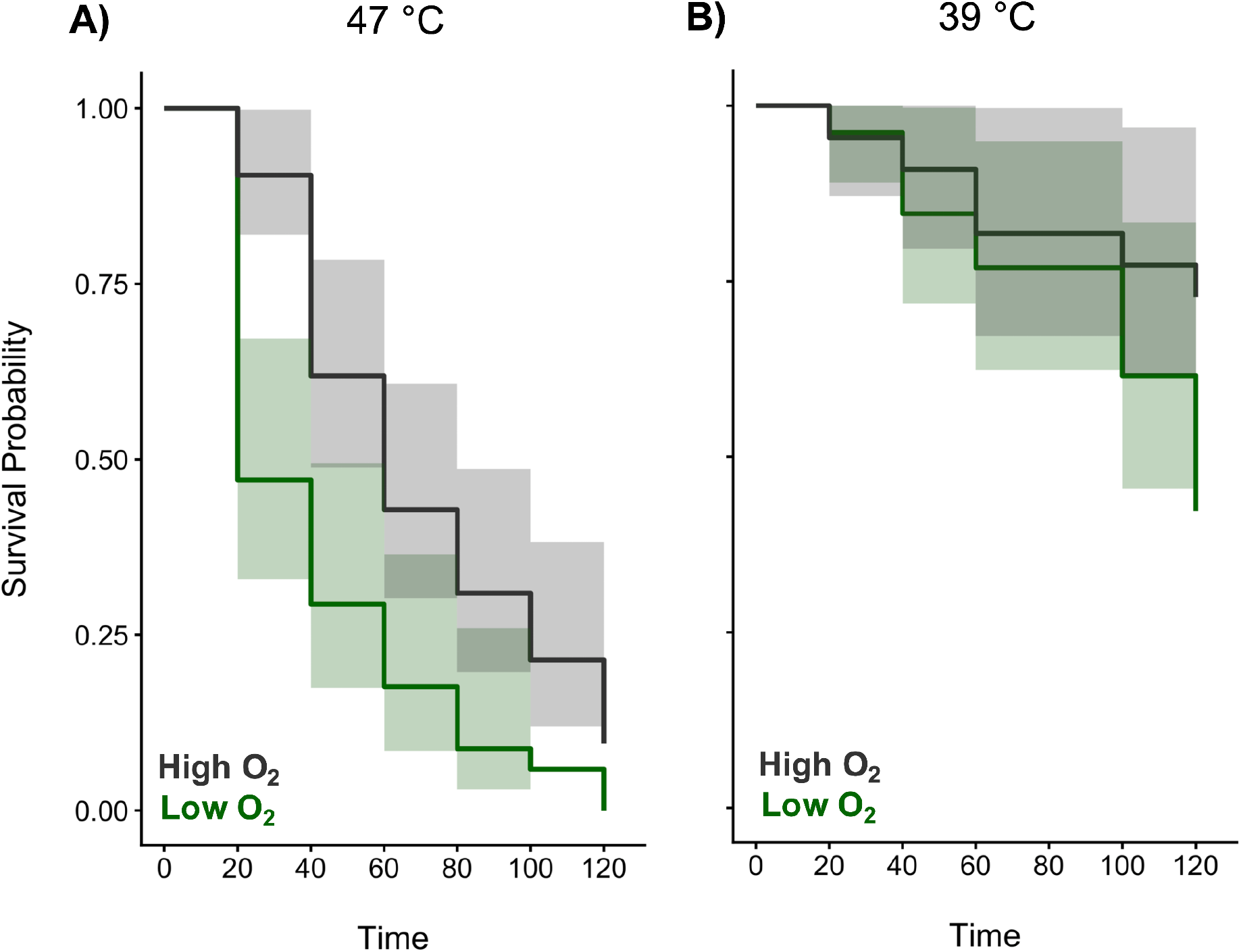
Low oxygen and high temperatures reduce survival of adult dragonflies. A) Estimated survival curves of adults tested at 47ºC. C) Estimated survival curves of adults tested at 39ºC. Adults came from *P. longipennis, L. pulchella*, and *E. simplicicollis*. Lines are those fitted from Cox-Proportional Hazards Models±95CIs.

### Does a species’ sensitivity to low oxygen conditions shape its upper elevational limit?

Among the 9 species in our phylogenetic generalized least squares, the species that are capable of living at higher elevations in Colorado also had lower mortality rates under low oxygen (parameter estimate±SE=-0.25±0.07, 95%CIs: -0.39 to -0.12; Fig. 2). This pattern is even stronger when we subset the analysis to include only the 7 species for which we have at least 4 observations for mortality rate in the focal treatment combination (parameter estimate±SE=-0.34±0.04, 95%CIs: -0.42 to -0.26).

**Fig. 2.**
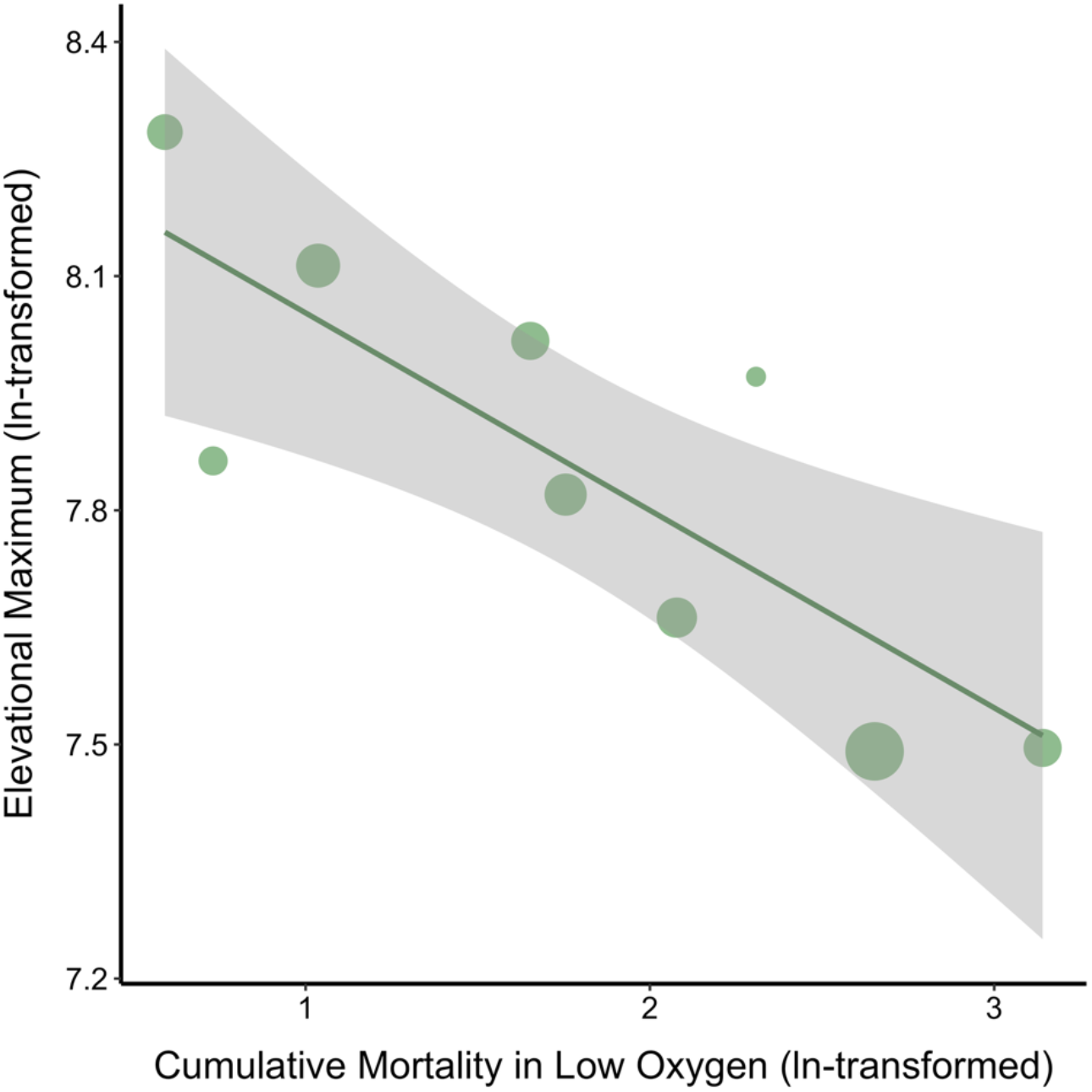
Species that live higher in the mountains of Colorado are also more tolerant of low O_2_. Points are estimated-marginal mean mortality values for each species as well as its elevational maximum in Colorado, USA. Line is best-fitted relationship between mortality in low O_2_ and elevational maximum from a phylogenetic generalized least squares analysis ±95%CIs. Size of points is proportional to the sample size.

## DISCUSSION

Low oxygen is one of the most notable challenges of life in the mountains [1–3], yet very little is known about its impact on the upper elevational limits of terrestrial ectotherms [3]. Contrary to arguments that altitudinal hypoxia is not a major issue for air-breathing insects and other ectothermic species, our results show that the oxygen at high elevations (∼4100m asl) is a serious threat to adult dragonflies (Fig. 1). Moreover, we also found that the species that can inhabit the highest elevations are also the most tolerant of low oxygen (Fig. 2). Collectively, these results provide the first direct evidence that hypoxia could set species’ elevational limits in a group of terrestrial ectotherms.

To date, the vast majority of prior research on the elevational limits of ectotherms has focused on the constraints imposed by the cold or, sometimes, food availability [1,2,30]. Our findings here show that low oxygen is a lethal problem even for immobilized dragonflies, and species’ elevational limits are strongly predicted by how well they tolerate low oxygen. A prior biogeographic analysis of dragonflies found that species that live at higher elevations across North America also tend to have biomechanical adaptations that lower the aerobic requirements for flight [22]. Transplant experiments remain necessary, but the emerging picture is that hypoxia could help set the elevational limits of dragonflies. Of course, hypoxia is probably not the only physiological problem for dragonflies at high elevations [22,31]. Indeed, that same biogeographic analysis also showed that adaptation to the cold was still the strongest predictor of species’ elevational maxima [22]. Our results therefore argue that low oxygen should be considered among the several major issues that dragonflies must overcome to live at high elevations.

Earlier work had argued that the insect tracheal system is so efficient that insects can deliver enough oxygen to meet resting demands even at Earth’s highest elevations [6,9]. Instead, we found that immobilized dragonflies are killed by the oxygen that naturally occurs just barely above the highest known records of dragonfly populations in the Rocky Mountains [26]. It seems then that hypoxia could be a major barrier to montane life among dragonflies, rejecting the notion that low oxygen is never a problem for terrestrial insects or other air-breathing ectotherms. Nevertheless, it is reasonable to wonder if our results generalize to other animals. Dragonflies do have extraordinary demands for oxygen [20], and a recent study using the same experimental design even showed that an elevationally migrating moth (*Euxoa auxiliaris*) was not at all sensitive to the same low-oxygen conditions that we used here [11]. Perhaps, then, dragonflies are unusual. Before making that conclusion, however, it is important to note that no other comparative studies have yet tested if physiological differences in oxygen tolerance predict the upper elevational limits of terrestrial ectotherm species [3]. Since our findings indicate that low oxygen can limit the elevational maxima of species with very high oxygen demands, it would be valuable to perform more studies about altitudinal hypoxia in ectotherms with lower oxygen demands.

While our results show that low oxygen is a serious threat to dragonflies, even this assessment might still be a significant underestimate for a few reasons. First, we examined the impacts of low oxygen on immobilized dragonflies. Nevertheless, in order to persist at high elevations or any other environment, dragonflies must hunt for prey, avoid predators, defend territories, mate, and engage in many other behaviors [32]. Critically, oxygen demands can be >30 times greater when flying insects, like dragonflies, are active compared to when they are at rest [6,33]. High-elevation dragonflies may therefore face even more dire oxygen shortfalls than were observed here. Second, our results demonstrate a potential constraint on air-breathing adults, yet the threat of hypoxia is thought to be an even more severe issue in the aquatic environments where dragonflies spend their larval stage [13,34]. Biophysical analyses and prior empirical work have shown that aquatic stages can be quite vulnerable to functional hypoxia at high elevations, leading to behavioral changes and fitness reductions in both the short and long term [3,6,13,35,36]. Thus, even though we found that low oxygen is a serious threat to adults, the adult stage might not be part of the life cycle that is actually the most sensitive to this environmental challenge. Clearly, future work assessing the effects of high-elevation conditions across both an animal’s daily routine and its life cycle will be necessary to understand the true impact of low oxygen on species’ persistence at high elevations [4,37,38].

Despite collecting all of the dragonflies from the same low-elevation sites, species differed in how vulnerable they were to low oxygen. It remains to be seen what drives these interspecific differences, but one possibility is variation in the number, size, and location of their tracheae [2,30,39,40]. Indeed, prior work has showed that some insects respond to hypoxia by increasing the total tracheal volume [39,40]. Tracheation like this is stimulated by hypoxia-inducible factor (HIF) signaling, which may also differ across species [6]. Another possibility is variation in the activity of key enzymes used in oxidative phosphorylation, like cytochrome c oxidase (COX). Prior work shows that high-elevation Tibetan locusts have evolved enhanced cytochrome c oxidase activity, which contributes to mitochondrial robustness and maintains high aerobic respiration in their mountain habitats [18]. A similar alteration in COX has also been shown in high-altitude bar-headed geese [41], and activity of the COX pathway was higher for leaf beetles at high than low elevations [15]. Though future work will need to evaluate the underlying proximate mechanisms, our findings reveal that even species from the same habitat can exhibit differences in oxygen tolerance that may have facilitated range expansion into higher elevations.

As we project into the future, species are facing novel threats due to warming conditions and increases in extreme heat events at lower elevations [42,43]. Indeed, our results show that hot conditions are also a significant threat to adult dragonflies. Heat at lower elevations is currently pushing many animal species upslope where thermal conditions are more tolerable [2,44,45]. However, our results in dragonflies also indicate that low oxygen could set limits that prohibits species from dispersing too far into the high elevations. If other species are also sensitive to low oxygen [3], then many animals could face a dilemma: the risk of overheating pushes the trailing edge of species’ ranges upwards, while the risk of hypoxia prevents the leading edge of species’ ranges from moving much at all. Indeed, a recent analysis of elevational range shifts across >800 insect species suggests that this hypothesized phenomenon is already occurring for many groups [17]. It is hard to predict the outcomes of such a range contraction for dragonflies or any other taxa, but it is likely to bring more species into contact and set the stage for competition between species in novel contexts [2,46,47]. While there is potential for animals to overcome these barriers via behavioral or plastic and/or evolutionary modifications [48], it is increasingly doubtful that species can outpace the speed of anthropogenic change [49]. Thus, for species that cannot handle the low oxygen at high elevations, our results suggest that life in the mountains will become much harder in the coming years.

## Supporting information

Supplemental Methods and Results

## Acknowledgements

We thank M. Young and M.-L. Le for help in the field. A. Lenard, N. Leith, and J. Stroud provided valuable feedback on the project. MPM, ARB, and MJM were supported by NSF Award #2532582.

