## Supplemental Methods and Results for "Hypoxia can set the elevational limits of dragonflies"

### SUPPORTING METHODS & RESULTS

*Can low oxygen be a problem for dragonflies at rest?*

We placed dragonflies in a DigiTherm Incubator set to a hot but sub-lethal temperature (47ºC; [23]) and measured how long each adult survived under either an ambient oxygen treatment for our study location ($P_{O2}$=130.7mm Hg) or a low-oxygen treatment ($P_{O2}$ = 95.7mm Hg). To manipulate the oxygen levels inside the incubator, we used a Hypoxico® Sierra 100 altitude generator, which filters $O_2$ out of the ambient air and flows a specified gas mixture into a space without altering overall air density. We flowed air into the incubator at 100 mL/min by inserting the altitude generator's outflow hose into a hole that was drilled through the sidewall of the incubator [11]. The chosen oxygen levels correspond to the levels found around Denver ($P_{O2}$ = 130.7mm Hg at ~1600 m asl) and around the elevation just above the highest known occurrences of dragonflies in Colorado (e.g. $P_{O2}$ = 95.7mm Hg at ~4,100 m asl). All manipulations were performed in the same incubator on different days. We performed one experimental treatment per day, and so all dragonflies that were captured on the same day were exposed to the same conditions (see also [11] for validation of these methods).

To compare patterns of mortality between the ambient and low-oxygen treatments, we fit mixed-effects Cox Proportional Hazards Models using the R package 'coxme' [25]. Adults that died during the treatment were scored as undergoing the “event” (event=1), with their death time as the time of their last known fate. Adults that did not die during their trial were scored as not having an event (event=0), with 120 minutes as the time of their final known fate. In this model, we included a fixed effect for the oxygen treatment. We also fit models where mass was included as a continuous fixed effect, but this term did not improve models, and so it was not considered further. To account for any non-independence of adults that were from different species, or tested in the same treatment but on different days, we included random intercepts for species and date. Overall, this analysis included 109 individuals from 9 species. For this analysis, we report the fold-change in mortality rate ± SE induced by low oxygen and evaluate the hypothesis by evaluating if the 95%CIs of the parameter estimate overlap 1 (i.e. 1x = no change). If low oxygen is a lethal problem for adult dragonflies, then the low-oxygen conditions should induce a fold-change in mortality rate that is greater than 1.

*Can low oxygen be a problem for dragonflies at rest?*

Dragonflies subjected to low oxygen had 3.1x ± 0.3 higher mortality rates than those subjected to ambient oxygen (95%CIs: 1.6x to 6.0x).

**Table S1.** $AIC_c$ table comparing models of mortality under different oxygen and temperature levels. Evidence ratios report how much more likely the top-model is compared to the focal model.

| **Terms** | **$AIC_c$** | **Akaike Weights** | **Evidence Ratio** |
|---|---|---|---|
| O2 + Temp + O2:Temp | 702.1993 | 0.357 | 1.7 |
| O2 + Temp | 701.1125 | 0.615 | 1 |
| O2 | 707.7336 | 0.022 | 27.4 |
| Temp | 710.9730 | 0.004 | 138.4 |
| Null | 716.4026 | 0.000 | 2090.3 |

**Table S2.** AICc model comparison for the best fitting model of phylogenetic non-independence in the comparative analysis of species' upper elevational limits and their oxygen tolerance. df

| Evolutionary Model | df | AICc |
|---|---|---|
| Brownian Motion | 3 | 22.584 |
| Pagel's Lambda | 4 | 14.956 |
| Star Phylogeny | 3 | 9.878 |
